# Exploring alternative quorum sensing model structures and quorum quenching strategies

**DOI:** 10.1101/2023.07.07.548074

**Authors:** Chiara Cimolato, Massimo Bellato, Gianluca Selvaggio, Luca Marchetti, Giulia Giordano, Luca Schenato

## Abstract

Bacterial quorum sensing (QS) is a cell-to-cell communication mechanism through which bacteria share information about cell density, and tune gene expression accordingly. Pathogens exploit QS to orchestrate virulence and regulate the expression of genes related to antimicrobial resistance. Despite the vast literature on QS, the properties of the underlying molecular network are not entirely clear. We compare two synthetic QS circuit architectures: in the first, a single positive feedback loop autoinduces the synthesis of the signal molecule; the second includes an additional positive feedback loop enhancing the synthesis of the signal molecule receptor. Our comprehensive analysis of the two systems and their equilibria highlights the differences in the bistable and hysteretic behaviors of the alternative QS structures. Finally, we investigate three different QS inhibition approaches; numerical analysis predicts their effect on the steady-state behavior of the two different QS models, revealing critical parameter thresholds that guarantee an effective QS suppression.

## Introduction

Bacterial cells are able to interact in various forms; cooperation enables them to sense and respond to their environment and coordinate collective behaviors thanks to several sophisticated signaling systems [1, 2].

The most widely known form of cooperative interaction among bacteria is *quorum sensing* (QS), a cell-to-cell communication system by which bacteria modify the expression of specific genes in relation to cell population density, through a positive feedback regulation involving the production, release, and detection of small signaling molecules. Even though bacterial species differ in QS-associated genes and signal molecules, the overall mechanism is the same: when the sensed bacterial population density is low, each cell produces low basal-levels of signal molecules, which are also released out of the cell and gradually diffuse into the environment. As the cell population increases, the signal molecules accumulate inside and outside the cells. Inside the bacteria, these signaling molecules bind to a transcriptional regulator (*receptor*), and the resulting active complex induces the transcription of specific QS-regulated genes. In particular, the production of signal molecule synthases is up-regulated, which gives rise to a positive feedback loop that autoinduces the synthesis of more signaling molecules (which are therefore called *autoinducers*).

This peculiar autoinducing mechanism is of particular interest in both microbiology and synthetic biology. In fact, the production of virulence factors and the activation of mechanisms that allow antimicrobial resistance (AMR), such as biofilm formation and enhanced horizontal transfer, are among the activated phenotypes triggered by QS at high cell density [3]. On the other hand, QS has also been exploited by synthetic biologists for the dynamical population-level control of bacteria and for the potential development of innovative therapies [4]. The control of bacterial populations can range from cell density control [5] to dynamic metabolic engineering [6] and modulation of physiological processes [7]. In this context, QS has been successfully combined with other genetic circuits so as to synthesize synchronized transcriptional toggle switches [8], genetic oscillators [9] and genetic logic gates [10].

Both when considering QS mechanisms orchestrated by pathogens and when designing ad-hoc QS-based circuits in synthetic biology, the functioning of QS can be difficult to interpret and contrasting conclusions are reported in the literature [11, 12]. Mathematical models and systems-and-control approaches are precious to enable a deeper understanding of the QS machinery through both theoretic analysis and computational studies, and also to design control strategies that either enhance quorum sensing (when it is desired to perform a prescribed functionality) or suppress it (to hinder the activity of pathogens). The pioneering work in [13–15] laid the foundation for the development of mathematical models of quorum sensing. In the wake of these early models, QS has been modelled resorting to several different approaches: from deterministic differential equation models, which describe the dynamics of the concentrations of the involved key players based on rate equations [16–23], to stochastic models that capture the intrinsic stochasticity underlying molecular reactions [24–27], also including diffusion mechanisms and spatial constraints [28–31]. While existing models have successfully depicted many aspects of the bacterial QS system and have supported the design of synthetic genetic circuits exploiting QS with sufficient predictability, full agreement on the precise molecular network underlying QS has not been reached yet, especially for some bacterial species for which it is still challenging to gain an accurate understanding of the molecular mechanisms that govern QS [12, 18].

Moreover, resistant bacteria are an increasingly recognized threat [32]. AMR caused 60% of hospital infections already in 2002 [33], is now among the leading causes of death worldwide, with 1.27 million deaths attributed to AMR in 2019 [34], and is projected to cause 10 million deaths each year by 2050 [34, 35]. In this context, it is crucial to explore alternative treatments for bacterial infections [36] and QS is a possible target [33]. In particular, quorum quenching (QQ), namely the inhibition of quorum sensing mechanism, has been proposed as promising approach for therapeutic strategies, with studies aimed to discover QS inhibitors and design QQ synthetic circuits to treat bacterial infections by reducing antimicrobial resistance activation [37, 38].

In this work, we gain insight into the remarkable cooperative abilities of bacteria and the structures that underlie quorum sensing interactions based on the well-known LuxI/LuxR QS system (hereinafter referred to as the Lux system), which is active in the whole family of Gram-negative bacteria and is natively responsible for the regulation of bioluminescence in the bacterium *Vibrio fischeri* [39]. Despite the lower clinical relevance of the Lux system, plenty of experimental data are already available in the literature and thus allow us to perform simulations with relevant parameters so as to explore general features of different QS interaction patterns, rather than focusing on a species-specific QS system. Specifically, we conduct an analysis of two synthetic Lux-based QS circuits, where the underlying molecular network includes either one or two feedback interactions, and we compare their equilibria and the asymptotic behavior of the system trajectories. For both QS structures, our analysis reveals a bistable behaviour, where crossing a threshold in the number of cells drives the system from a low stable equilibrium (associated with the OFF state of the related genes, at low cell numbers) to a high stable equilibrium (associated with the ON state and with the activation of AMR-related genes, at high cell numbers); the presence of hysteresis is shown to confer robustness to the QS mechanism, once it has been activated, with respect to disturbances and fluctuations in the number of cells. Finally, we simulate and compare the steady-state behavior of the two synthetic systems in the presence of three different QS inhibition strategies. In particular, QS mechanisms can be inhibited by degrading extracellular autoinducers, or by sequestering the receptors to which autoinducers bind, or by reducing the production of autoinducer synthases. The goal is to destroy bistability, and therefore hinder the activation of the QS mechanism and of AMR-related genes in pathogens, by making sure that the system exclusively admits the low equilibrium. To this aim, we identified critical parameter thresholds that prevent the activation of QS communication, which are crucial to devise and experimentally design QQ strategies.

## Results

### Two Quorum Sensing circuit architectures

Gram-negative bacteria typically rely on homologous QS systems that are composed of three main elements:

- a gene *G*_1_ encoding for a synthase *S*, which is an enzyme responsible for the synthesis of a specific signaling molecule *A*, called autoinducer;
- a gene *G*_2_ encoding for a receptor, or transcriptional regulator, *R*, which is a protein that binds to the autoinducer to create an active complex *C*;
- a regulated promoter *P*, inducible by the active complex *C*, which drives the expression of the synthase *S*.

The uptake of QS signaling molecules has two effects on the cell. First, these signaling molecules control a range of activities directly related to antimicrobial resistance, pathogenicity, and biofilm formation. Second, these signaling molecules increase their own production, and are thereby called autoinducers. The QS mechanism is assumed to enable the expression of some population phenotypes only when a sufficient cell density is reached, while such cooperative behaviors are not cost-effective when the bacterial population density is too low. This behavior is the result of the positive autoinductive loop, as is visualised in Figure 1. As the amount of synthase *S* increases, the synthesis of the signaling molecule *A* also increases. This upregulates the concentration of the receptor-autoinducer complex *C*, which then results in an even larger increase in the production of synthase *S*. Notably, the same complex *C* also drives the transcription of a plurality of genes that are directly involved in antimicrobial resistance, pathogenicity, and biofilm formation: a strong correlation has been observed between quorum sensing activation and the increase of bacterial virulence and resistance to therapies [40].

**Fig. 1:**
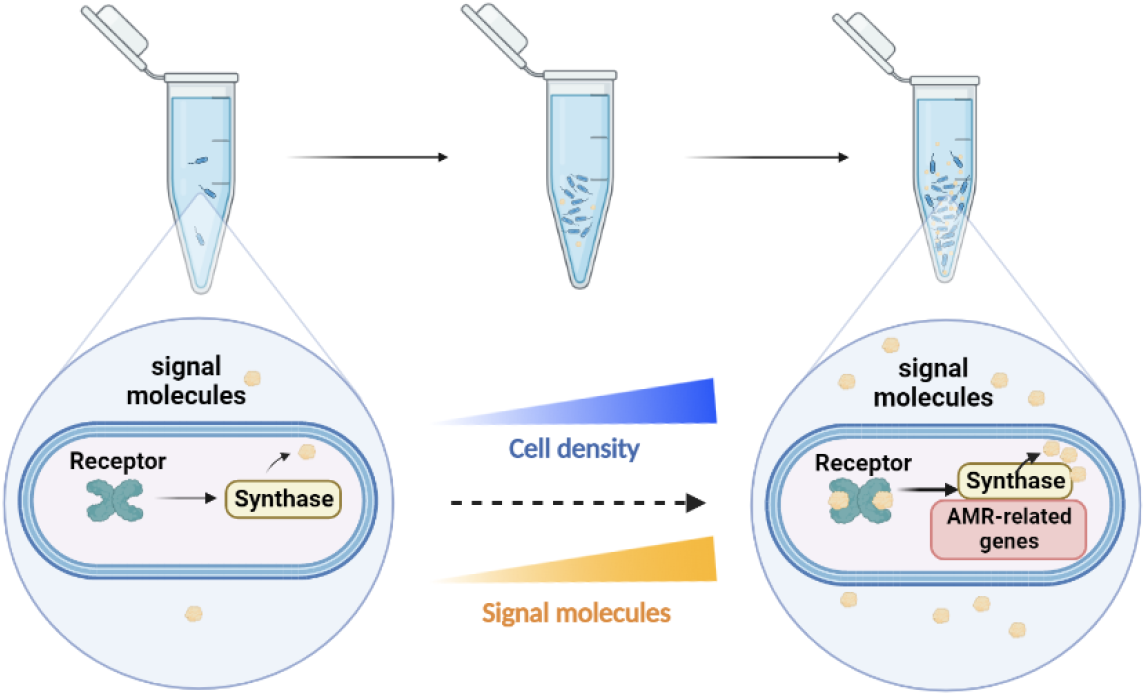
Quorum sensing mechanism: the higher the cell density, the higher the concentration of signaling molecules in each cell, which corresponds to an increased expression both of the synthase itself and of AMR-related genes in bacteria.

The phenomenon of quorum sensing was first observed in *Vibrio fischeri*, a bioluminescent marine bacterium, where QS controls light emission by regulating the production of luciferase enzymes [41]. In particular, once bacterial density crosses a certain threshold, the genes encoding for luciferase enzymes switch from the OFF to ON state. We take the QS network that describes these underlying processes, called Lux system, as an example of prototypical QS mechanism. A schematic of the genetic circuit observed in *V. fischeri* is shown in Fig. 2. The Lux system consists of:

**Fig. 2:**
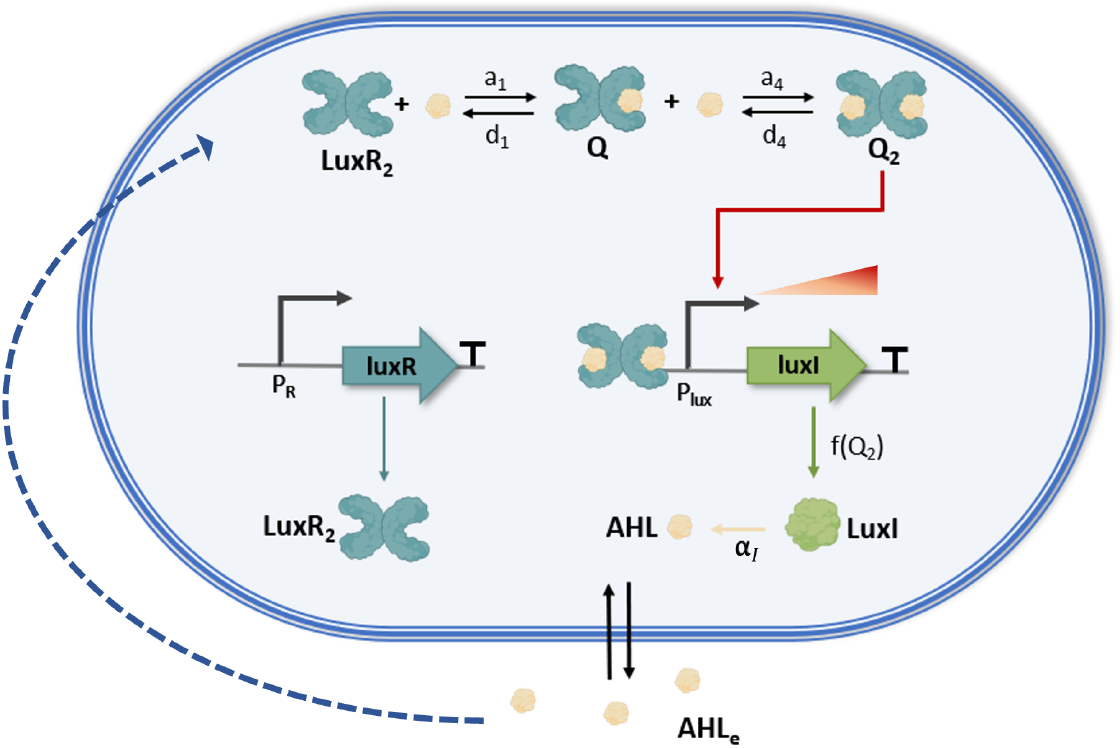
Schematic of the quorum sensing network *Lux system*, with a single positive feedback loop. The activated complex consists of a hetero-tetrameric structure of two LuxR molecules bound with two molecules of AHL [42].

- the synthase LuxI (encoded by the gene *luxI*), an enzyme which synthesizes a small signaling molecule called acyl-homoserine-lactone (AHL);
- the autoinducer AHL, which rapidly accumulates in and out the cell, diffusing outside the cell through the membrane;
- the LuxR protein, encoded by the gene *luxR*, whose expression is constitutively regulated by the promoter P_R_.

When the LuxR protein dimerizes, it is able to detect and bind to the intracellular autoinducer AHL, thus forming the LuxR-AHL active complex. The active complex then induces the regulated promoter P_lux_ by binding with the *lux box* promoter site, and thus regulating the expression of *luxI* and closing the positive feedback loop. Therefore, once the bacterial density reaches a critical threshold level, the binding of the active complex with the promoter increases the transcription rate of LuxI from its basal to its maximum level.

### Single-feedback Lux system

We develop a dynamical mathematical model that represents an engineered bacterial QS network reproducing the Lux system with a single feedback loop. This synthetic QS Lux system, implemented in host strains of the bacterium *Escherichia coli* [42, 43] and shown in Fig. 2, can be described by the ordinary differential equation model

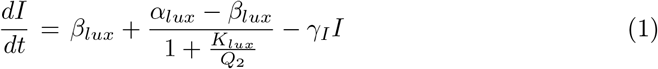

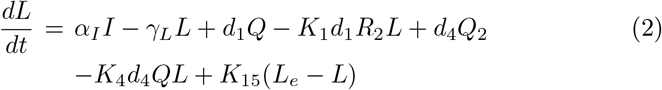

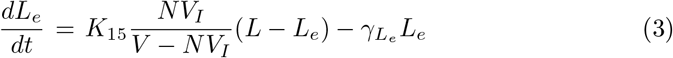

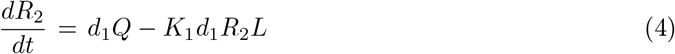

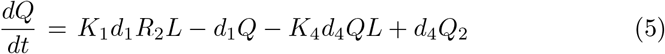

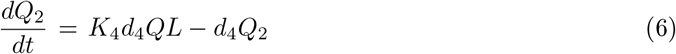

where *I* represents the concentration of protein LuxI (arbitrary units per cell), *L* and *L*_*e*_ respectively denote the intracellular and extracellular concentrations of AHL (in nM), *R*_2_ indicates the concentration of LuxR homodimer (in nM), while *Q* and *Q*_2_ denote the concentrations (in nM) of LuxR homodimer bound to one molecule of AHL (intermediate complex) or two molecules of AHL (active complex), respectively. Numerical values of model parameters were taken from [42–48].

Equation (1) describes the time evolution of the concentration of LuxI includes two synthesis terms: *β*_*lux*_ is the basal synthesis rate in the OFF state, while the inducible synthesis term depends on *Q*_2_ as a Hill function, where *α*_*lux*_ is the maximum synthesis rate per cell and *K*_*lux*_ corresponds to the AHL concentration to reach the half-maximum inducible synthesis rate. The last term in (1) represents the spontaneous degradation of LuxI with rate *γ*_*I*_.

In the equation (2) ruling the evolution of intracellular AHL concentration, *α*_*I*_ is the rate at which AHL is synthesized by LuxI, *γ*_*L*_ the spontaneous degradation rate of AHL. As shown in Fig. 2, *a*_1_ and *d*_1_ are the forward and backward rate constants for the binding and unbinding reactions of the first AHL molecule to the LuxR homodimer, respectively. The equilibrium constant for this reaction is defined as *K*_1_ = *a*_1_*/d*_1_, and therefore in equation (2), according to mass action kinetics, we include the consumption term *a*_1_*R*_2_*L* = *K*_1_*d*_1_*R*_2_*L* together with the production term *d*_1_*Q*. The forward and backward rates describing the second AHL binding reaction are *a*_4_ and *d*_4_, respectively, with equilibrium constant *K*_4_ = *a*_4_*/d*_4_, which leads to the consumption term *a*_4_*QL* = *K*_4_*d*_4_*QL* and the production term *d*_4_*Q*_2_ in equation (2). The last term in equation (2) accounts for the diffusion process of the autoinducer across the membrane, with *K*_15_ being the AHL permeability rate of the wall of *E. coli* cells [45].

Equation (3) describes the dynamics of the concentration of extracellular AHL, where *N* is the number of spatially homogeneous bacteria in the culture, *V*_*I*_ is the volume of a single *E. coli* cell and *V* is the volume of the entire cell culture environment; moreover, the degradation rate of *L*_*e*_ is 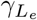.

Equation (4), capturing the time evolution of the concentration of LuxR homodimer, only includes the terms related to the unbinding and binding of LuxR homodimer to the first AHL molecule (with rates *d*_1_ and *a*_1_ = *K*_1_*d*_1_, respectively); synthesis and degradation terms of LuxR are omitted, since the protein is assumed to be constitutively produced at a constant rate that exactly compensates for its degradation.

Equation (5) describes the dynamics of the concentration of the intermediate complex (LuxR homodimer bound to one molecule of AHL) and only includes the terms related to the unbinding and binding of LuX homodimer to the first AHL molecule (with rates *d*_1_ and *a*_1_ = *K*_1_*d*_1_, respectively) and to the unbinding and binding of the intermediate complex to the second AHL molecule (with rates *d*_4_ and *a*_4_ = *K*_4_*d*_4_, respectively). Equivalently, equation (6), capturing the dynamics of the concentration of the active complex (LuxR homodimer bound to two molecules of AHL), only includes the terms related to the unbinding and binding of the intermediate complex to the second AHL molecule (with rates *d*_4_ and *a*_4_ = *K*_4_*d*_4_, respectively). The degradation terms for both *Q* and *Q*_2_ are neglected, since the binding of AHL to the LuxR homodimer is known to stabilize the homodimer [49].

We can observe that 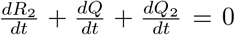, meaning that the sum of the three species concentrations is constant at all times. Introducing the constant 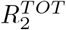, this leads to the following mass conservation law:

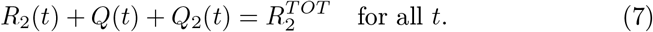

Moreover, since intracellular AHL can exist either in a free form (*L*), or bound to the LuxR homodimer in one copy (in *Q*) or in two copies (in *Q*_2_), we can introduce the total quantity

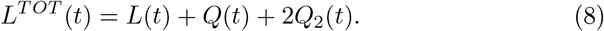

We assume that the probability of AHL binding to free sites is identical for *R*_2_ and *Q*, and the probability of AHL unbinding from occupied sites is identical for *Q* and *Q*_2_, which couples the parameter values through the following constraints:

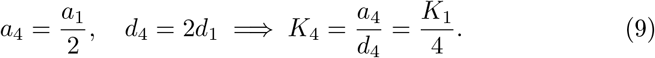

### Single-feedback Lux system: reduced-order model and equilibria

We investigate the steady-state behavior of the QS system (1)-(6) within a spatially homogeneous bacterial population. We rely on a quasi-steady state approximation, based on the assumption that the reversible reactions forming complexes *Q* and *Q*_2_ have fast dynamics and therefore *Q* and *Q*_2_ immediately reach their steady-state values, so that their derivative becomes zero. Timescale separation arguments allow us to obtain their equilibrium concentrations by setting equations (5) and (6) to zero. Then, recalling that 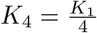, we have

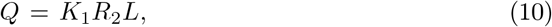

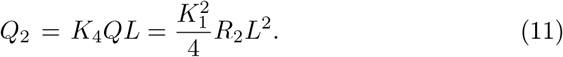

Exploiting the conservation law in equation (7) leads to the following steady-state concentrations for *R*_2_, *Q* and *Q*_2_:

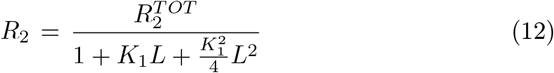

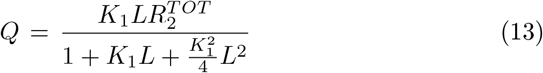

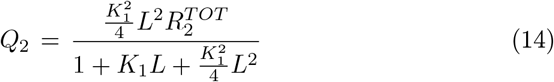

As a consequence, the six-variable model (1)-(6) can be reduced to the three-variable system:

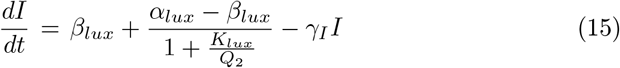

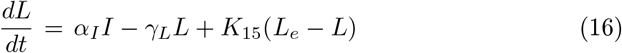

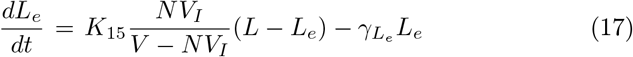

where *Q*_2_ has the constant value in equation (14).

We can now compute the equilibria of the reduced-order system, by setting the equations (15)-(17) to 0. By defining 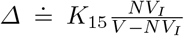, some algebraic manipulations yield the linear equilibrium conditions

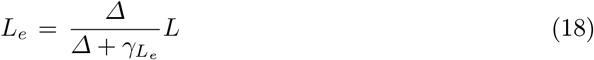

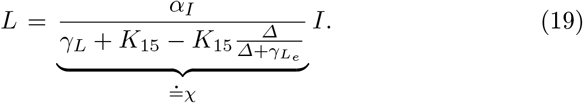

Namely,

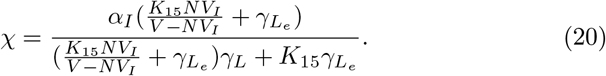

Now, we can rewrite *Q*_2_ as a function of *I* by substituting equation (19) into equation (14), and we get

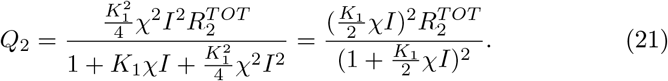

By substituting equation (21) into equation (15), the steady-state value of *I* can be obtained by solving equation

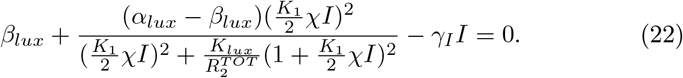

Fig. 3 visualises the equilibrium points (steady-state concentrations *Ī* of LuxI per cell), for different values of the cell density *ρ* = *N/V*, as the inter-sections of the curves 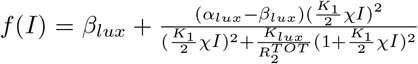, associated with LuxI synthesis, and *g*(*I*) = *γ*_*I*_*I*, associated with LuxI degradation. Different values of *ρ* lead to different behaviors, ranging from monostability (at either low or high values) to bistability.

**Fig. 3:**
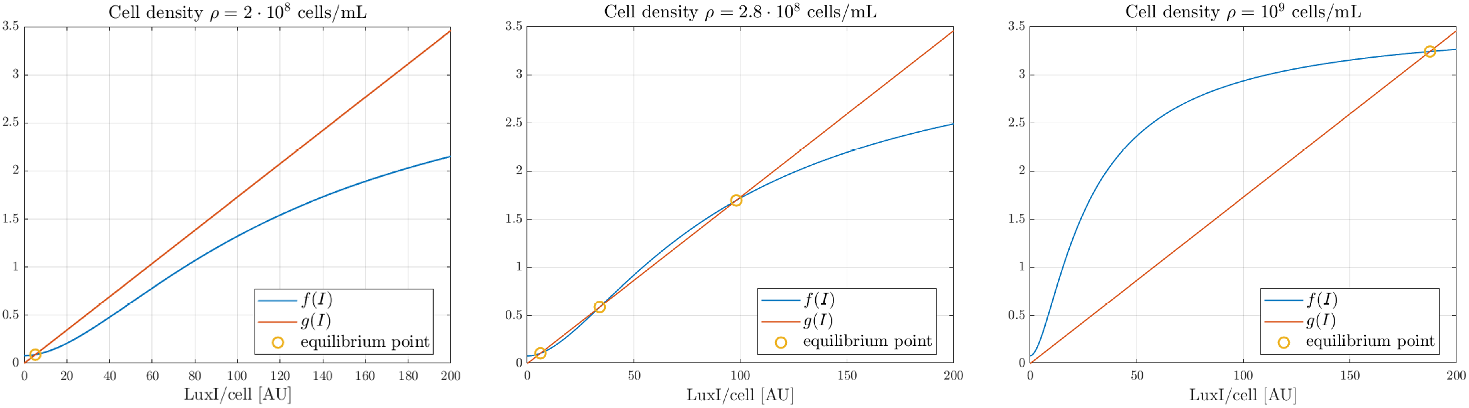
LuxI equilibrium points *Ī* in the single-feedback model for different values of the cell density *ρ* = *N/V*. The steady-state concentrations of LuxI per cell correspond to the intersections of *f* (*I*), associated with LuxI synthesis, and *g*(*I*), associated with LuxI degradation. In the left panel, for *ρ* = 2 · 10^8^ cells/mL, the system admits a unique low stable equilibrium (OFF state). In the central panel, for *ρ* = 2.8 10^8^ cells/mL, the system admits three equilibria (two stable, for low and high values, and one unstable, for intermediate values). In the right panel, for *ρ* = 10^9^ cells/mL, the system admits a unique high stable equilibrium (ON state).

Through the coefficient *χ* = *χ*(*N*), the system equilibria strongly depend on the number *N* of cells inside the culture environment. For this reason, we compute the solution *Ī* = *Ī* (*N*) of equation (22) for different values of *N*. In fact, since the culture volume is fixed (and assumed equal to *V* = 1 mL) and the cell density can be expressed as [cells*/*mL], varying the number of cells *N* is equivalent to varying the cell density *ρ* = *N/V*. Fig. 4 shows the bifurcation diagram of the steady-state concentration *Ī* of LuxI per cell against the population density *ρ*. For low values of the cell density, the system admits a unique equilibrium, corresponding to the basal concentration of LuxI in the OFF state, which is stable. When the population density increases above a certain threshold, the system starts exhibiting a bistable behavior, because at the same cell density three equilibria coexist: two stable equilibria (a high stable equilibrium and a low stable equilibrium) along with an intermediate unstable equilibrium. If the cell density further increases above a second threshold, the system again admits a unique equilibrium, associated with a high concentration of LuxI per cell (ON state), which is stable.

**Fig. 4:**
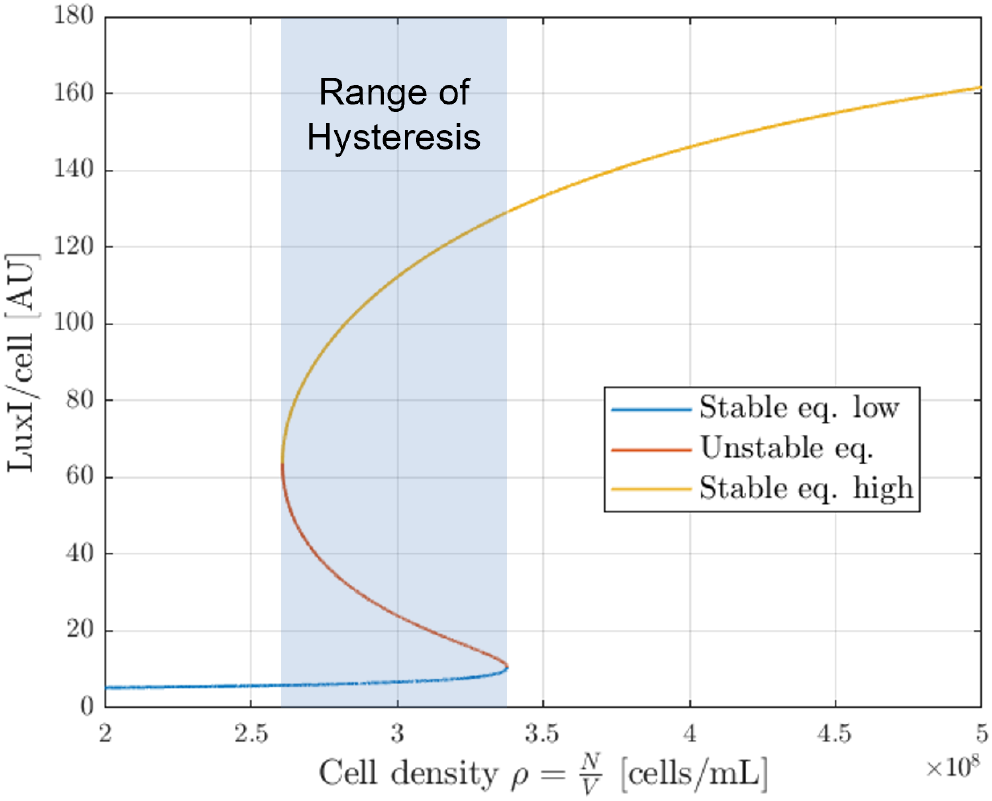
Bifurcation diagram of the steady-state concentration *Ī* of LuxI per cell for varying cell density *ρ*.

As can be seen from Fig. 4, the bacterial colony exhibits a hysteretic response: the cell density necessary to activate the communication process is higher than the cell density needed to inactivate it, thus making the communication process more robust to perturbations in the cell density.

### Double-feedback Lux system: LuxR up-regulation

The cell-to-cell communication system based on the Lux mechanism has been well characterized experimentally. However, models alternative to the one we have analyzed in the previous section have been proposed in the literature. In particular, evidence has been provided in [33] for the existence of a second feedback look that concurs to the formation of the QS network in some bacterial families. In the double-feedback Lux system, shown in Fig. 5, the active complex, whose concentration we previously denoted as *Q*_2_, increases the synthesis not only of the synthase enzyme LuxI (with concentration *I*), but also of the receptor LuxR (with concentration *R*_2_), presumably by binding to a second *lux-box* promoter site. Therefore, similarly to the autoinducer AHL, the receptor LuxR enhances its own synthesis depending on the sensed cell density.

**Fig. 5:**
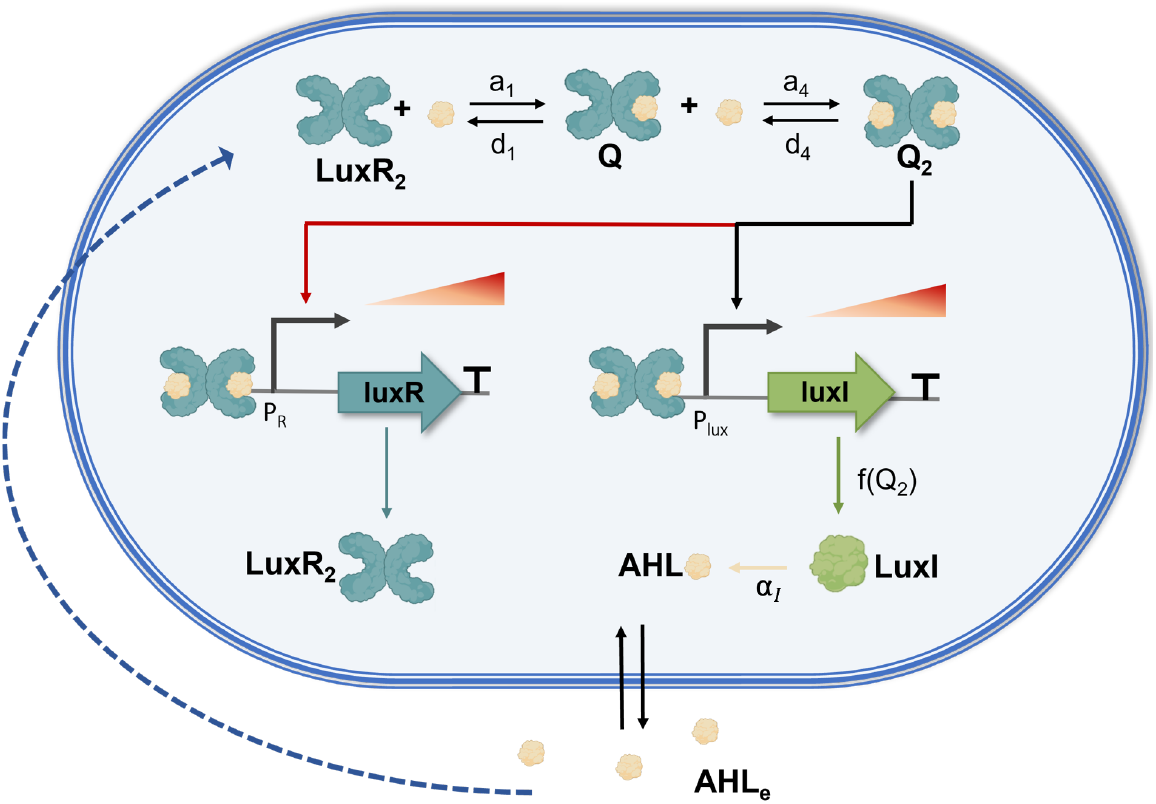
Schematic of the quorum sensing network *Lux system* with two positive feedback loops.

There is no consensus in the scientific literature on the number of feedback loops that are present in QS systems. Some authors claim that the receptor is only synthesized at a constant basal level, as in our model (1)-(6): most of the QS mathematical models that have been proposed, see e.g. [28, 50], only include the autoinduction loop affecting the signaling molecule. Other works, conversely, propose models that also take into account a second feedback loop acting on the receptor [12, 16, 17, 51].

Here, we propose a mathematical model of an engineered bacterial QS network that reproduces the Lux system with both feedback loops, and we explore its equilibrium properties to gain insight into the circuit behavior. This synthetic QS Lux system that includes both positive feedback loops, implemented in host strains of the bacterium *E. coli* and visualized in Fig. 5, can be described by the ordinary differential equation model

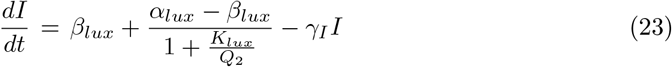

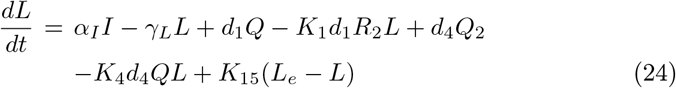

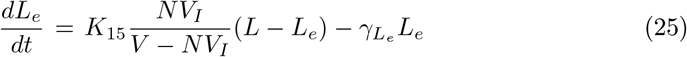

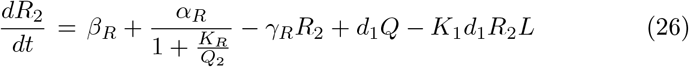

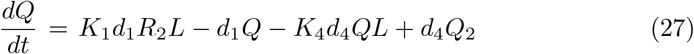

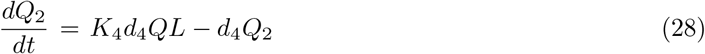

The only difference with respect to the single-feedback model (1)-(6) is in equation (26), which, in addition to the terms already present in (4), also includes spontaneous degradation with rate *γ*_*R*_ as well as LuxR homodimer synthesis, expressed by two terms: *β*_*R*_ is the basal synthesis rate of *R*_2_ when the gene is in the OFF state, while the inducible synthesis term depends on *Q*_2_ as a Hill function, where *α*_*R*_ is the maximum inducible synthesis rate and *K*_*R*_ corresponds to the AHL concentration that yields the half-maximum inducible synthesis rate.

Now 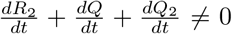, hence we have no mass conservation. Still, we can observe that the LuxR homodimer can exist either in a free form (*R*_2_), or bound to one copy of intracellular AHL (in *Q*), or bound to two copies of intracellular AHL (in *Q*_2_), and therefore we can introduce the time-varying total quantity

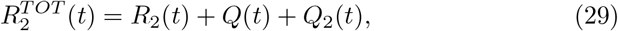

while the total quantity of intracellular AHL still evolves over time as

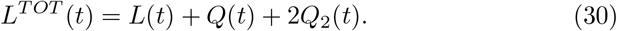

Again, we can assume that variables *Q* and *Q*_2_ reach their steady state much faster than the others (because the corresponding species are involved in fast reactions only). In view of timescale separation arguments, we can obtain their equilibrium concentrations by setting equations (27) and (28) to 0. This yields *Q* = *K*_1_ *R*_2_ *L* and 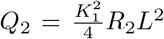, as in the single-feedback QS model. Then, to find the equilibria of the double-feedback QS model, we set equations (23)-(26) to 0. Since the first three equations are unchanged with respect to the previous model, we can exploit (19) to rewrite *Q*_2_, now as a function of both *I* and *R*_2_:

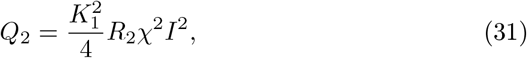

where *χ* is defined in (20). Using the above relation, the system equilibria can be obtained by solving the system of equations

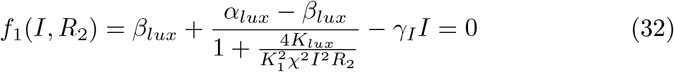

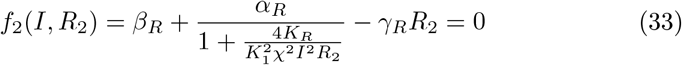

The intersections (*Ī*, 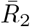) of the nullclines (32) and (33) correspond to the system equilibria, as shown in Fig. 6 for different values of the cell density *ρ* = *N/V*. The equilibrium landscape strongly depends on parameter *N*, the number of cells within the culture environment. Fig. 7 shows the bifurcation diagram of the steady-state concentration *Ī* of LuxI per cell, for varying cell density *ρ* = *N/V*, for the double-feedback model and also for the single-feedback model.

**Fig. 6:**
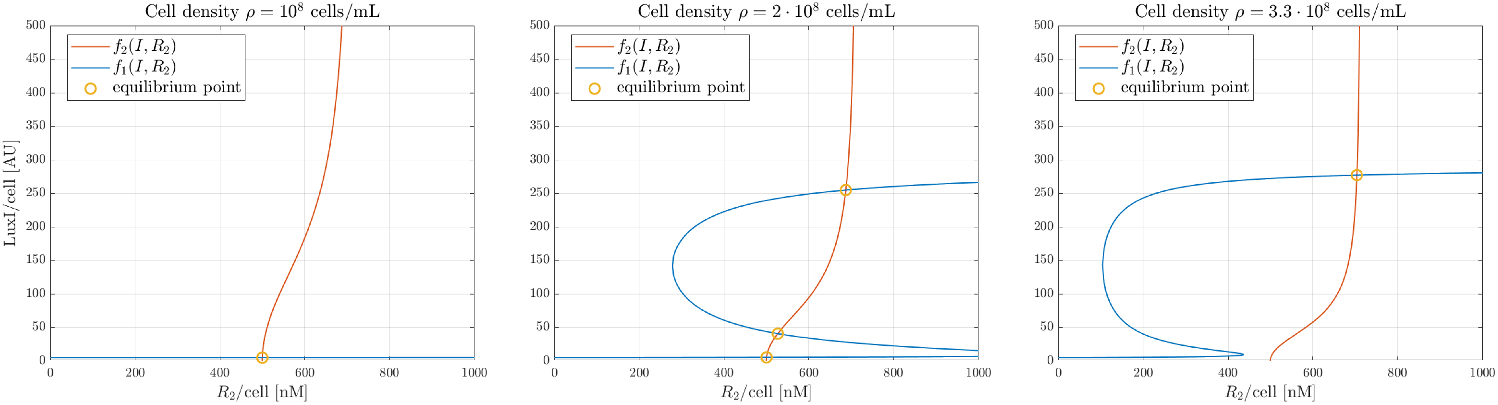
Equilibrium points (*Ī*, 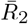) in the double-feedback model for different values of the cell density *ρ* = *N/V*. The steady-state concentrations of *R*_2_ and LuxI per cell correspond to the intersections of the nullclines *f*_1_(*I, R*_2_) and *f*_2_(*I, R*_2_). In the left panel, for *ρ* = 10^8^ cells/mL, the system admits a unique stable equilibrium, associated with low values of LuxI (OFF state). In the central panel, for *ρ* = 2 10^8^ cells/mL, the system admits three equilibria (two stable, for low and high values of LuxI, and one unstable, for intermediate values of LuxI). In the right panel, for *ρ* = 3.3 10^9^ cells/mL, the system admits a unique stable equilibrium, associated with high values of LuxI (ON state).

**Fig. 7:**
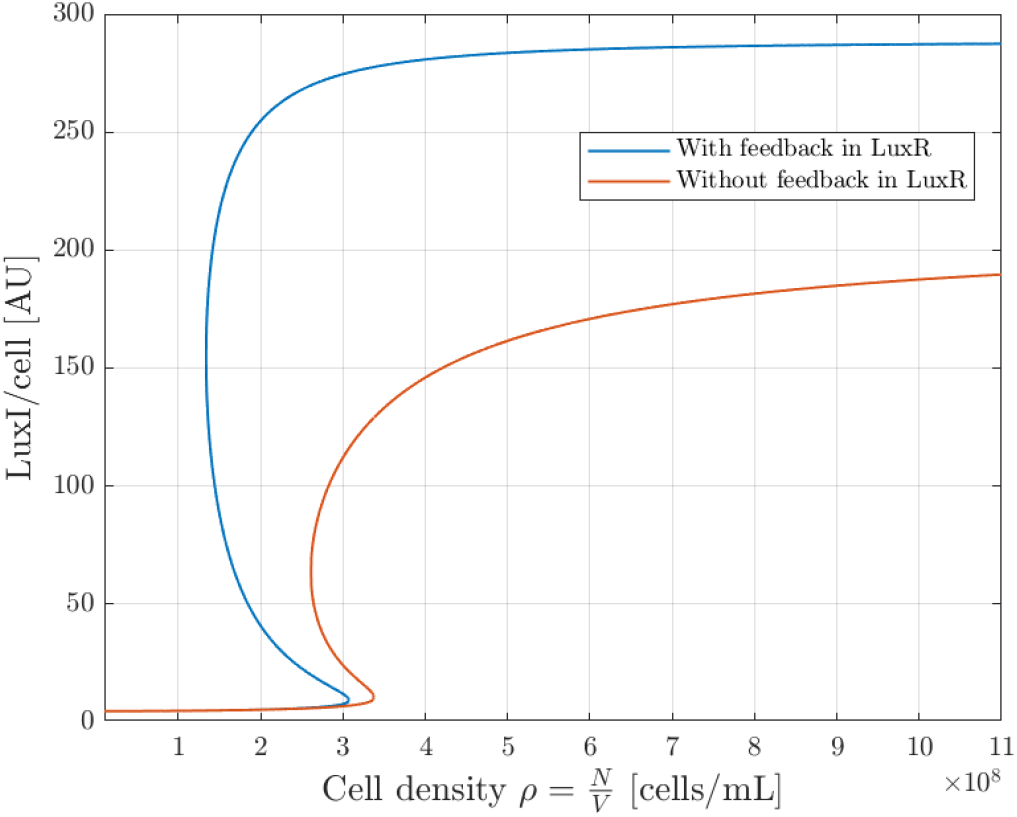
Bifurcation diagram of the steady-state concentration *Ī* of LuxI per cell for varying cell density *ρ*, for the double-feedback Lux system (blue) and for the single-feedback Lux system (red). The presence of the second positive feedback loop increases both the range of hysteresis and the value of the high stable equilibrium.

By comparing the bifurcation diagrams for the two models, we can observe that the presence of the second positive feedback definitely increases the range of hysteresis: in fact, the first threshold occurs at much smaller values of cell density, even though the second threshold occurs at slightly smaller values. A broader hysteresis range notably improves the robustness of the system with respect to fluctuations in the cell density. Moreover, the maximum steady-state concentration of LuxI per cell is much larger in the double-feedback system, leading to a much higher stable equilibrium in the ON state. As already mentioned, the QS mechanism is activated much earlier (the threshold occurs at much smaller cell density values) in the double-feedback system.

The bifurcation diagram shown in Fig. 7 demonstrates a strong correlation between the size of the cell population and the QS mechanism switching from the OFF to the ON state. Since the metabolic and resource-related cost for the activation of the QS communication process is not negligible, the QS mechanism is activated only when the population size is sufficiently large to create a net population benefit (namely, above the first threshold); then, once QS has been activated, hysteresis confers robustness with respect to cell density variations, thus preventing accidental inactivation of the QS mechanism due to a temporary fluctuation in the number of cells.

### Modeling QQ and QS inhibition strategies via key parameter tuning

To contrast the ever-increasing health threat posed by antimicrobial resistant bacteria, novel therapeutic techniques are needed. In view of the involvement of the QS mechanism in providing antibiotic resistance to bacteria, inhibiting bacterial QS is a promising strategy to control bacterial infections [33]. To disrupt the QS communication process, QS inhibitors and Quorum Quenching enzymes have been discovered [52], which can contribute to restoring the susceptibility of bacteria to antibiotics.

QS inhibitors are designed to interfere with the QS molecular communication system in bacterial pathogens, thus preventing cells from detecting neighboring cells and therefore reducing the expression of QS-regulated genes, which govern virulence factors and biofilm formation, among the causes of chronic diseases.

Several sites of the QS pathway can be targeted to interfere with the cell-to-cell communication system. Here we explore three different strategies:

i. degrading extracellular AHL molecules, thanks to degrading enzymes that prevent extracellular AHL accumulation [43];
ii. sequestering AHL-receptor proteins via antagonist compounds [53];
iii. reducing the production of the synthase, exploiting techniques that allow for sequence-specific regulation of gene expression (e.g., interference repressors based on CRISPRi and RNAi).

We now assess the potential effectiveness of the three therapeutic strategies applied to our two different models. In each model, the various strategies are implemented by altering specific model parameters, so as to simulate the treatment effect.

In particular,

i. the enzyme-driven degradation of extracellular AHL can be represented by increasing the degradation rate 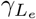 in equation (3) for the first model and in equation (26) for the second model;
ii. since the sequestration of LuxR homodimers (AHL receptors) antagonizes their interaction with AHL to form complexes *Q* and *Q*_2_, it can be represented by a reduction in the binding rates *a*_1_ and *a*_4_ or, equivalently, in the equilibrium constants *K*_1_ and *K*_4_, in both models;
iii. the reduction of the synthesis of LuxI can be represented by a smaller maximum LuxI synthesis rate per cell *α*_*lux*_ in equation (1) for the first model and in equation (23) for the second model.

The corresponding results are reported in Fig. 8, where the impact of the three inhibition mechanisms described above is simulated by considering both the single-feedback and the double-feedback Lux QS systems.

**Fig. 8:**
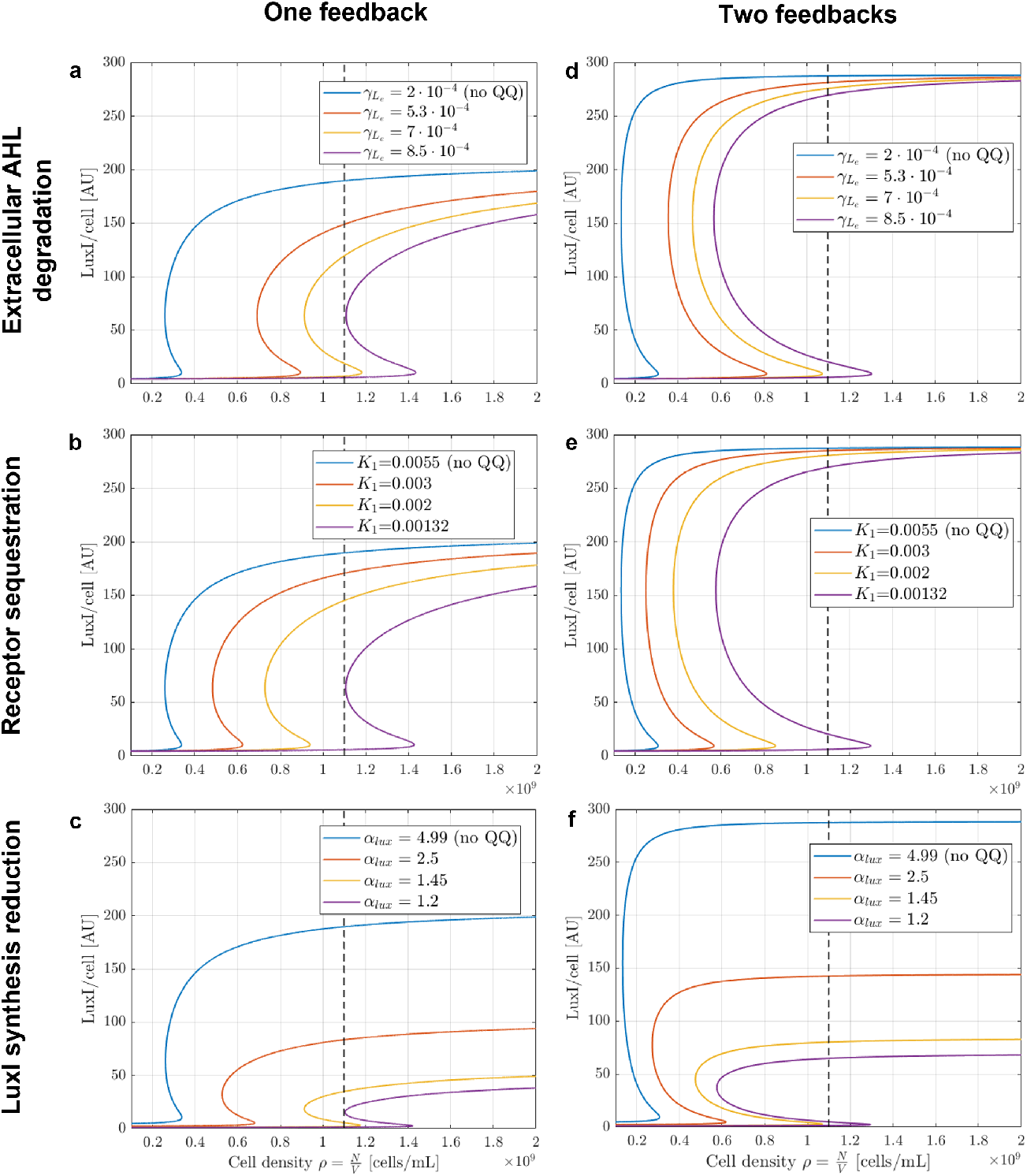
Effects on the LuxI bifurcation diagram of three different QS inhibition strategies: extracellular AHL degradation (panels a and d), receptor sequestration (panels b and e), LuxI synthesis reduction (panels c and f). In each plot, the simulations are performed for increasing values of the inhibition strength. Dashed vertical lines correspond to the maximum cell density achievable for *E. coli* in a culture environment. In panels a, b and c the simulation results are obtained by considering the dynamics of the single-feedback Lux QS model, while the double-feedback Lux QS model dynamics are considered in panels d, e and f.

Previous studies have revealed that the upper limit on the cell density of *E. coli* in a culture environment is approximately 1.1 *·* 10^9^ cells/mL [47, 54], which is shown as a dashed vertical line in the plots of Fig. 8. Consequently, the disruption of bacterial communication is guaranteed provided that QS molecules production remain at the basal (low) level until the maximum density is reached.

In all the simulations of Fig. 8, the bifurcation diagram is depicted in blue when no inhibition is present, while in other colours (red, yellow, purple) when the inhibition strategy is implemented with increasing strength.

The left column of Fig. 8 shows the simulations associated with the three QS inhibition strategies applied to the single-feedback QS system, for varying values of the parameters associated with the corresponding inhibition strategy. The bifurcation diagrams showing the concentration of LuxI per cell, as a function of cell density, for different values of 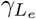, *K*_1_ and *α*_*lux*_ are illustrated in panels a, b and c respectively. A strong enough inhibition can ensure that the system is monostable: it admits a unique equilibrium, which is stable, associated with low values of LuxI concentration, for all achievable cell densities. In particular, for all the three different strategies applied to the single-feedback model (left column of Fig. 8), the parameter values corresponding to the purple curves lead to a sufficiently strong inhibition of QS: at all feasible cell densities, the system has a unique stable equilibrium, which corresponds to the OFF state of the cell-to-cell communication process; therefore, the bistable region can never be reached. The simulations, based on our previous equilibrium analysis, allow us to identify the key threshold values of 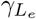, *K*_1,4_ and *α*_*lux*_ that prevent the activation of QS, even at high cell densities: for the single-feedback system, the inhibition of QS is guaranteed when 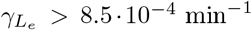 *>* 8.5 *·* 10^*−*4^ min^*−*1^ (the degradation of extracellular AHL is sufficiently increased by the presence of degrading enzymes), or *K*_1_ *<* 1.3210^*−*3^ nM^*−*1^ (the AHL-receptor binding rate is sufficiently reduced by the presence of antagonistic sequestering compounds), or *α*_*lux*_ *<* 1.2 AU min^*−*1^ (the production of the synthase is sufficiently reduced by interference repressors).

The right column of Fig. 8 shows the simulations that illustrate the effect of the three QS inhibition strategies on the LuxI bifurcation diagrams for the double-feedback QS system. The same values of 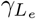, *K*_1_ and *α*_*lux*_ as in the left column are considered. Consistently with our previous analysis, we see that the double-feedback system is more robust and therefore, in this case, inhibiting the cell-to-cell communication system requires a stronger action. Also, the QS mechanism is activated at much lower cell densities, which makes inhibition even more challenging. It is worth noting that synthase synthesis reduction technique has a twofold effect: it not only increases the cell density threshold at which the communication system switches on, but also reduces the maximum LuxI steady state concentration.

## Discussion

Quorum sensing is a prominent example, among several regulatory pathways, showing the ubiquity of control strategies in natural systems: QS mechanisms allow bacteria to perform robust disturbance rejection by employing feedback control techniques. As the amount of quantitative data concerning QS molecular networks increases, mathematical models have become crucial to unravel and understand the complex structure of these networks and their feedback architecture.

Different QS molecular network structures have been identified and suggested in the literature. To compare them, we have proposed two different models of QS architectures implemented through an engineered Lux QS circuit in homogeneous bacterial populations of *E. coli*. The single-feedback model is characterized by a single positive loop that enhances the production of autoinducer. The double-feedback model also includes a second positive loop for the autoinductive regulation of the receptor expression. Our formal analysis of equilibria and steady-state behaviors has revealed that both systems exhibit bistability for a suitable range of cell densities, along with an interesting hysteretic behavior that confers robustness and enables disturbance rejection once the QS mechanism has been activated. This leads to a memory effect in the circuit behavior, because the system response is not solely dependent on the current cell density, but also on its history. A comparison between the two model structures has shown that the double-feedback architecture exhibits increased robustness with respect to perturbations, in particular, to cell density variations: not only the QS communication mechanism is activated earlier, but also, once the mechanism is activated, inactivating it due to a reduction in cell density is much more difficult. While the bistable and hysteretic behavior of QS systems has already been identified in the literature [12, 17–19], these studies either considered a different type of underlying molecular network or used model parameters not derived from experimental data. After having gained more insight into the properties of these two QS motifs, we have investigated the effect of three different inhibition strategies aimed at hampering the QS communication system. The equilibrium analysis of the systems, with the parameters suitably modified by the three inhibition approaches, has identified key parameter thresholds to prevent the activation of the QS communication mechanism. Moreover, considering the same QQ parameter values, the inhibition strategies applied to the double-feedback system are less effective than when applied to the single-feedback system and, to obtain a similar result, higher inhibitions levels are needed.

These findings offer fundamental knowledge to predict and control Lux-like QS systems and represent an essential preliminary study to support the design of engineered biological systems for quorum sensing inhibition, or quorum quenching. In particular, our analysis has highlighted crucial experimental and architectural requirements for therapeutic approaches aimed at suppressing QS-based strategies that reinforce antimicrobial resistance in bacteria, through strategies that can be either pharmacological or based on synthetic biology.

In future studies, we aim to combine model predictions and high-resolution experiments, which will allow us to gain a better understanding of the underlying network (by identifying the most suitable model structure based on experimental data) and to explore new ways of controlling and inhibiting QS mechanisms.

## Methods

All the simulation code was implemented in Matlab^®^ 2021b (MathWorks, Inc). The solver ode15s was used to compute the system solutions with default option parameters. Equilibrium concentrations were computed using the symbolic solver vpasolve with default option parameters. The values of all the parameters, variables and constants adopted to numerically simulate the model equations are gathered from different articles that study and exploit the Lux system for both systems biology and synthetic biology applications [5, 42–44, 46–48].

## Acknowledgments

The authors thank Elisa Gaetan, Barbara Di Camillo, Simone Del Favero and Lorenzo Pasotti for helpful discussions.

## Funding

This work was funded by: the Fondazione Cariparo grant “Bando Ricerca Scientifica di Eccellenza 2021 n59576”; the University of Padova, Department of Information Engineering, PhD Grant PON-RI 2014-20 (CCI2014IT16M2OP005) on FSE REACT-EU funds; the University of Padova, Department of Information Engineering “type B Research Grant”.

## Authors’ contributions

CC, MB, GS, GG and LS conceived the study. CC and MB developed the models. CC performed the calculations and the numerical simulations, and prepared the figures. CC, MB and GG drafted the manuscript. GS, GG and LS supervised the work. LM and LS provided logistic and financial support. All the authors revised and approved the final manuscript.

## Notes

### Competing Interest Statement

The authors have declared no competing interest.

